# Expression of oxytocin receptors in the zebra finch brain during vocal development

**DOI:** 10.1101/739623

**Authors:** Matthew T. Davis, Kathleen E. Grogan, Donna L. Maney

## Abstract

Juvenile male zebra finches memorize and learn to sing the song of a male caregiver, or “tutor”, during a complex vocal learning process. Juveniles are highly motivated to interact socially with their tutor, and these interactions are required for effective vocal learning. It is currently unknown what neurological mechanisms underlie attraction to tutors, but social motivation and affiliation in this and other species may be mediated by oxytocin and related nonapeptides. Here, we used qPCR to quantify expression of oxytocin receptor (OTR) mRNA in the lateral septum, auditory forebrain, and regions of the song control system in zebra finches throughout post-hatch development and vocal learning. We found that zebra finches express OTR mRNA in these regions from post-hatch day 5 to adulthood, encompassing the entire period of auditory and sensorimotor learning. We also mapped the binding of ^125^I-ornithine vasotocin, an oxytocin receptor antagonist that binds to oxytocin receptors in songbird brain, to understand the neuroanatomical distribution of oxytocin-like action during vocal development. This study provides the groundwork for the use of zebra finches as a model for understanding the mechanisms underlying social motivation and its role in vocal development.

## Introduction

For juvenile songbirds, learning to sing is a highly social endeavor. Song is learned by copying a “tutor”, usually the father. The social influences on song learning have been best-studied in the zebra finch, which has become the most common songbird in laboratory research. In this species, only males sing, and song is learned during an early sensitive period after which exposure to new tutors does not result in new learning (reviewed by Gobes, 2019; see also Fig. 1). As juvenile male zebra finches approach adulthood and their songs become fully developed, those songs come to closely resemble the songs of the father. This effect is clearly not genetic – if a young male nestling is cross-fostered to a different set of parents, he will copy the song of the foster father even if he can hear his biological father singing in the same room. Remarkably, this effect has been observed even if the foster father is heterospecific (Immelmann 1969; Roper & Zann 2005). Thus, the social influences on song learning are quite robust and can even outweigh innate preferences for conspecific song. Further, live male tutors are more effective at promoting learning than playback of pre-recorded song (Derégnaucourt et al., 2013; Eales, 1989; Immelmann, 1969). Collectively, this evidence shows that social interactions provide potent learning signals that are required for normal vocal development. This species may therefore be a powerful model with which to understand the neural mechanisms underlying the development of social learning.

**Fig. 1.**
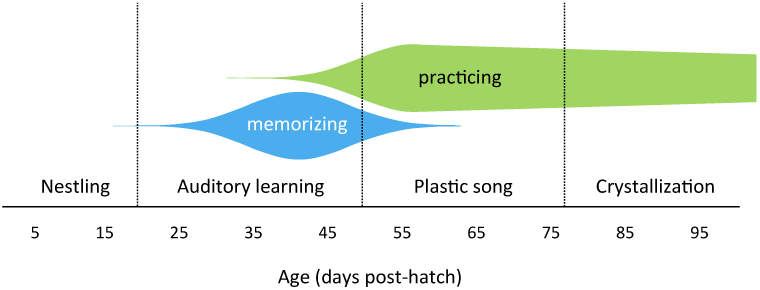
Developmental trajectory of song learning in zebra finches. Zebra finches fledge the nest at about post-hatch day 20 (P20); before this time, there is little evidence that they are able to form memories of tutor song (see Gobes et al., 2019 for review). By P25, the critical period of auditory learning has opened, and the young males begin to attend to and memorize male songs. They begin practicing song-like vocalizations, called subsong, a few days later (Arnold, 1975). By P35, song (particularly the tutor’s song) has acquired high incentive salience; young males will key-press to hear it (Tchernichovski et al., 1999). The incentive salience of tutor song peaks before P45; after P50, young males shift their auditory preference from tutor song to the songs of other males (C. A. Rodriguez-Saltos, et al., unpublished data). At about the same time, they enter the “plastic song” phase, during which their song rate increases dramatically (Johnson et al., 2002) and their song begins to resemble tutor song. After about P77, a male’s song is highly similar to its final form (Johnson et al., 2002), which will not change substantially in adulthood; it is becoming “crystallized”.

Given that song learning clearly has a social component, tutor choice may rely on the activity of neuropeptides that direct attention to social cues. Several independent groups of researchers, including ours, have hypothesized that attention and attraction to father’s song, which promote learning of that song, are mediated by oxytocin receptors (OTR) or related nonapeptide receptors (Baran, 2017; Maney & Rodriguez-Saltos, 2016; Theofanopoulou et al., 2017). OTR is thought to play a key role in the development of social attachment in a wide variety of vertebrate taxa (Gordon et al., 2011; Vaidyanathan & Hammock, 2017). Ross & Young (2009) demonstrated that in mice, administration of an oxytocin antagonist early in life resulted in reduced motivation of juveniles to return to caregivers after a period of separation. Similarly, Baran et al. (2016) showed that a vasopressin antagonist reduced affiliative interactions with caregivers in juvenile zebra finches. Thus, the role of nonapeptides in attraction to social cues, even during development, may be conserved across many species.

Birds have a clear homolog of mammalian OTR (Gubrij et al. 2005; Leung et al., 2011), which has been called the VT3 or “OT-like” receptor. Here, in keeping with a much-needed shift toward standardized nomenclature (Kelly & Goodson, 2014), we refer to it as OTR. In birds, the receptor has two endogenous ligands: mesotocin and vasotocin, which are the avian homologs of oxytocin and vasopressin, respectively. Both ligands are predicted to bind to OTR with high affinity (see Leung et al., 2011 for a discussion), and vasotocin has oxytocin-like functions in birds (Goodson & Bass, 2001; Robinzon et al., 1994; Saito & Koike, 1992). Avian OTR is expressed in many of the same brain regions as in mammals, including the lateral septum (Leung et al., 2009; 2011), in which OTR is thought to be important for social memory and social motivation (Goodson et al., 2009; Lukas et al., 2013). Local administration of oxytocin antagonist into this region reduces sociality and pair bonding in zebra finches (Goodson et al., 2009; Klatt & Goodson, 2013; Pedersen & Tomaszycki, 2012). In mice, OTR expressed in the auditory cortex may facilitate tuning of auditory neurons to conspecific vocalizations (Marlin et al., 2015); songbirds express OTR in auditory areas that respond preferentially to conspecific song (Leung et al., 2009; 2011). OTR is also expressed in HVC and dorsal arcopallium (Ad; Leung et al., 2009; 2011), both of which are key parts of neural circuits critical for song learning (Bottjer & Altenau, 2010; Doupe, 1993). These findings have formed the basis for our hypothesis that OTR mediates song learning by directing attention to tutors, facilitating the formation of social preferences, and gating auditory information to promote the development of selective encoding of tutor song (Maney & Rodriguez-Saltos, 2016).

A primary aim of this study was to map the neural distribution of OTR throughout the entire course of development in juvenile zebra finches. Maney and Rodriguez-Saltos (2016) presented preliminary evidence that OTR is expressed in HVC and the auditory forebrain in juveniles during song learning. Here, we present a more complete assessment of OTR expression, across the entire period of song learning, in both male and female zebra finches. We used autoradiography to map the neural distribution of binding of the oxytocin antagonist ^125^I-OVTA in both sexes at ten ages from post-hatch day 5 to 95 (adulthood). This approach allowed us to determine where OTR is likely to be expressed throughout the brain over the entire course of song learning. In addition, we performed a quantitative analysis of OTR mRNA expression at the same ages in four brain regions important for social motivation and/or song learning: the septal area (containing the lateral septum), the auditory forebrain, and two song-related areas, HVC and Ad. We predicted that expression of OTR would be higher in males than females, particularly in song control regions, during song learning.

## Results

In this study, we mapped the binding of oxytocin antagonist ^125^I-OVTA in the brains of juvenile zebra finches. This ligand was developed for use in rodent brain tissue (Elands et al., 1988) and has also been used to label OTR in several species of primate (reviewed by Freeman & Young, 2016). Although the ligand is highly selective for OTR in rodents, it was more recently shown to bind to both OTR and vasopressin 1a receptors (V1aR) in primates (see Freeman & Young 2016). Its relative affinity for avian nonapeptide receptors has not been thoroughly investigated (but see Leung et al., 2009; Ondrasek et al., 2018; Voorhuis et al., 1990). Songbirds have at least three different nonapeptide receptors that are expressed in brain: OTR, V1aR, and vasopressin 2 receptors (V2R) (Leung et al., 2011). In adults, the pattern of ^125^I-OVTA labeling strongly resembles that of OTR mRNA, not V1aR or V2R mRNA, in our regions of interest (Leung et al., 2011; Fig. S1). Therefore, we interpret ^125^I-OVTA labeling in these regions as predominantly binding OTR. The results of the autoradiography are shown in Figs. 2-5. We found that the pattern of ^125^I-OVTA labeling in juveniles largely resembles that of adults (Leung et al., 2009) even at the earliest phases of development we sampled. We focus below on regions that are related to our hypothesis that OTR activity promotes social attachment and song learning in juveniles.

**Fig. 2.**
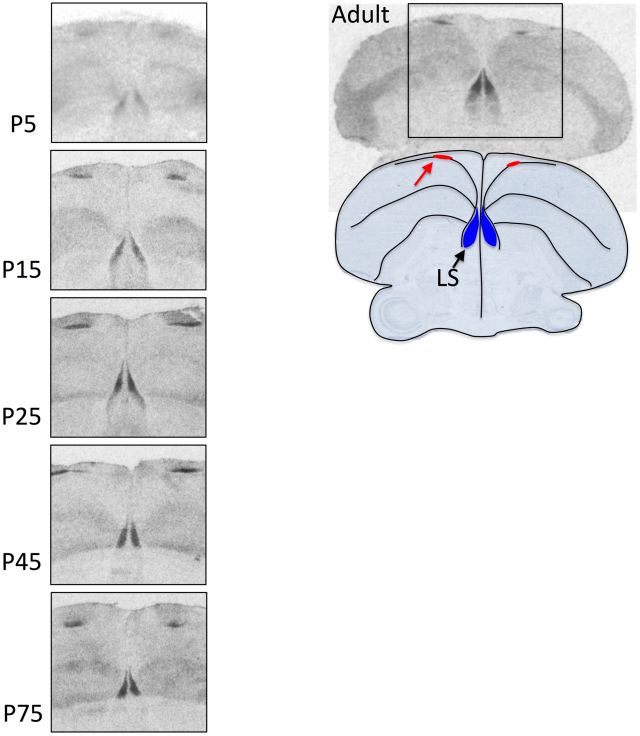
Binding of ^125^I-OVTA in the lateral septum. The upper right photo shows ^125^I-OVTA binding in an adult male zebra finch. Dense labeling can be seen in the lateral septum (LS) as well as in the periventricular area of the hyperpallium (red arrow in the drawing). On the left, the same labeling is shown at several ages post-hatch. The labeling was similar in males and females. All images are of coronal sections

**Fig. 3.**
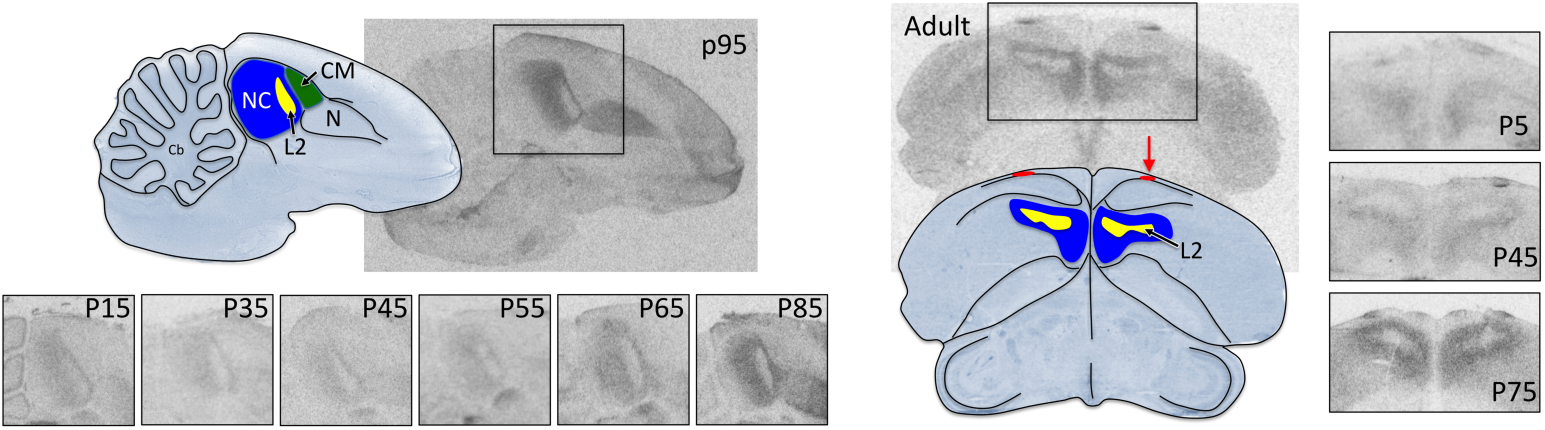
Binding of ^125^I-OVTA in the auditory forebrain. Labeling is shown in sagittal sections on the left, and coronal on the right. Dense labeling can be seen surrounding but not including Field L2 at all ages. Label is also notable in the periventricular area of the hyperpallium (red arrow). NC, caudal nidopallium. CM, causal mesopallium. N, nidopallium. Cb, cerebellum.

**Fig. 4.**
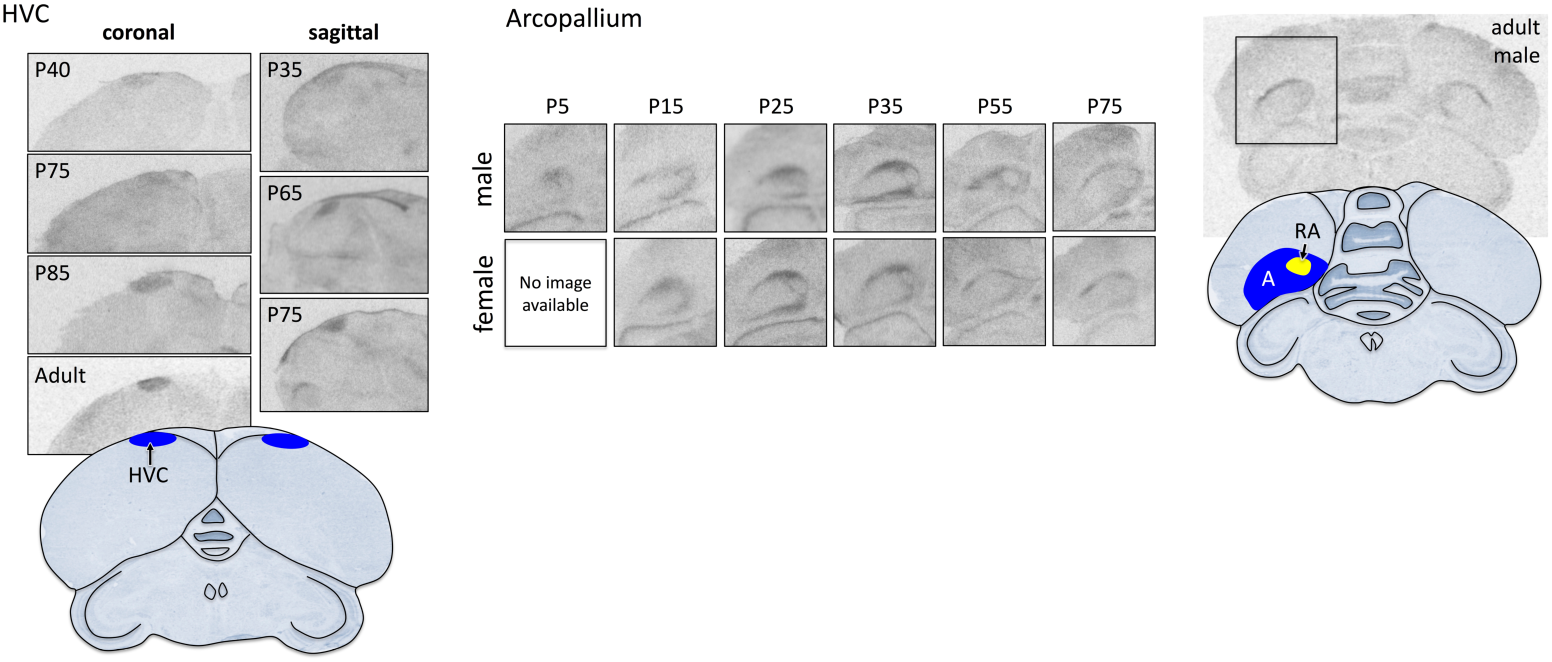
Binding of ^125^I-OVTA in HVC and the arcopallium. On the left, labeling is shown in song control nucleus HVC in male zebra finches over the course of vocal development. Coronal and sagittal views are shown; in the sagittal images, rostral is to the right. Above, labeling is shown in coronal images of the arcopallium (A) in both sexes. The arcopallium contains the robust nucleus of the arcopallium (RA), which is a major target of HVC. The dense label is seen in the dorsal arcopallium and the capsular region surrounding RA.

**Fig. 5.**
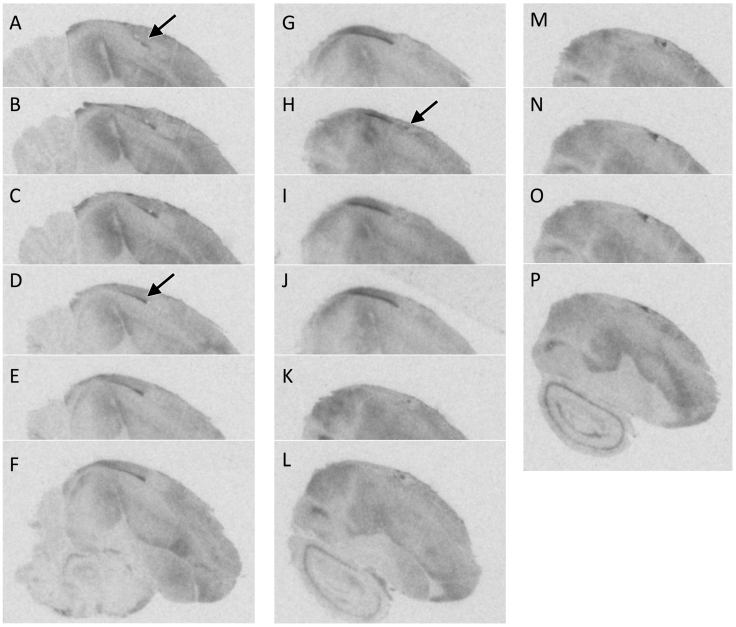
Binding of ^125^I-OVTA in the periventricular area of hyperpallium. This series of photos shows sagittal sections, lateralmost (A) to medialmost (P) in a female zebra finch at P85. Rostral is to the right. All sections are from the same individual and are 120μm apart. Labeling was similar at all ages in both sexes. Dense labeling can be seen in the periventricular area of the hyperpallium (arrow in A, becoming more obvious in D) that extends for several hundred μm medially, ending at J. A second, discontinuous area of label emerges at H and extends laterally to P.

Our qPCR analysis showed evidence of OTR expression in the four regions of interest throughout development. The developmental trajectories of expression are plotted for each sex in Fig. 6. Because nonapeptides bind somewhat promiscuously to OTR, V1aR, and V2R (reviewed by Kelly & Goodson, 2014), we also quantified expression of V1aR and V2R mRNA in these regions. We emphasize that OTR is likely the predominant receptor expressed in our regions of interest, however (Fig. S1). On average, expression of V1aR and V2R were low, with about 30% of the Cp values greater than 34. The data are presented in Figs. S2 and S3. Because we measured three different genes in four regions, we first performed an omnibus test (repeated-measures MANOVA, see Methods) that included all of the birds, regions and genes. This test showed main effects of region (p < 0.001) and gene (p < 0.001), as well as a region x gene interaction (p < 0.001), a region x gene x age interaction (p = 0.006), a region x gene x sex interaction (p = 0.001), and a region x gene x age x sex interaction (p = 0.020). We therefore proceeded to test for effects of age and sex within each gene and region. The results of those tests are presented in Table 1 for OTR, and Supplemental Table 1 for V1aR and V2R.

**Fig. 6.**
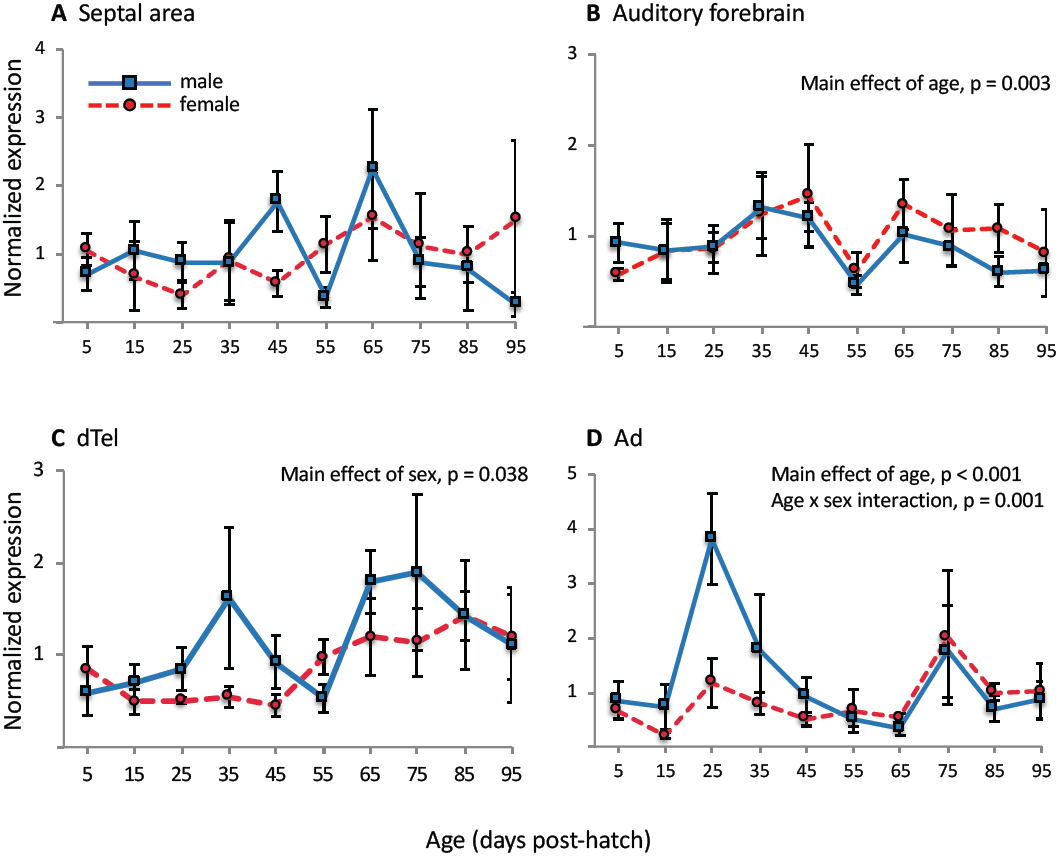
Developmental trajectories of oxytocin receptor mRNA expression in four brain regions of interest. Means and SEM are shown for males (blue) and females (red) at ten ages post-hatch. Expression was normalized to two reference genes for analysis, then normalized to the series mean (within region) for the purposes of graphing. Thus, 1.0 on the Y axis corresponds to the average across both sexes and all ages for each region, and region-to-region variation is not represented here. dTel, dorsal telencephalon (includes HVC when HVC is present). Ad, dorsal arcopallium.

**Table 1.** Results of univariate F-tests within gene and region, showing main effects of age (P5-P95) and sex on expression of oxytocin receptor mRNA, and interactions between age and sex, in the four regions of interest. Significant effects are shown in bold and marked by asterisks. Ad, dorsal arcopallium. dTel, dorsal telencephalon.

| Region | Age |  | Sex |  | Age x Sex |  |
| --- | --- | --- | --- | --- | --- | --- |
|  | F | p | F | p | F | p |
| Septum | 0.991 | 0.482 | 0.769 | 0.393 | 0.940 | 0.510 |
| Auditory forebrain | **4.441** | **0.003\*** | 1.980 | 0.176 | 1.076 | 0.423 |
| dTel | 1.977 | 0.112 | **8.078** | **0.012\*** | 1.394 | 0.269 |
| Ad | **17.509** | **<0.001\*** | 1.482 | 0.247 | **8.639** | **0.001\*** |

Because we had a maximum of only five individuals of each sex at each of the ten ages, we did not perform post-hoc pairwise tests for sex differences with each age. Instead, we grouped the birds into phases delineated by major developmental milestones (see Fig. 1), then compared the sexes within each phase. To guard against Type I error, we first ran an omnibus MANOVA, as above, with all three genes and four regions included, This analysis yielded a main effect of gene (p < 0.001), a main effect of region (p = 0.001), a region x gene interaction (p < 0.001), as well as a region x gene x sex x phase interaction (p = 0.037). We then tested for sex differences within gene and phase. The results of these tests are shown for OTR in Fig. 7 and for V1aR and V2R in Figs. S4 and S5.

**Fig. 7.**
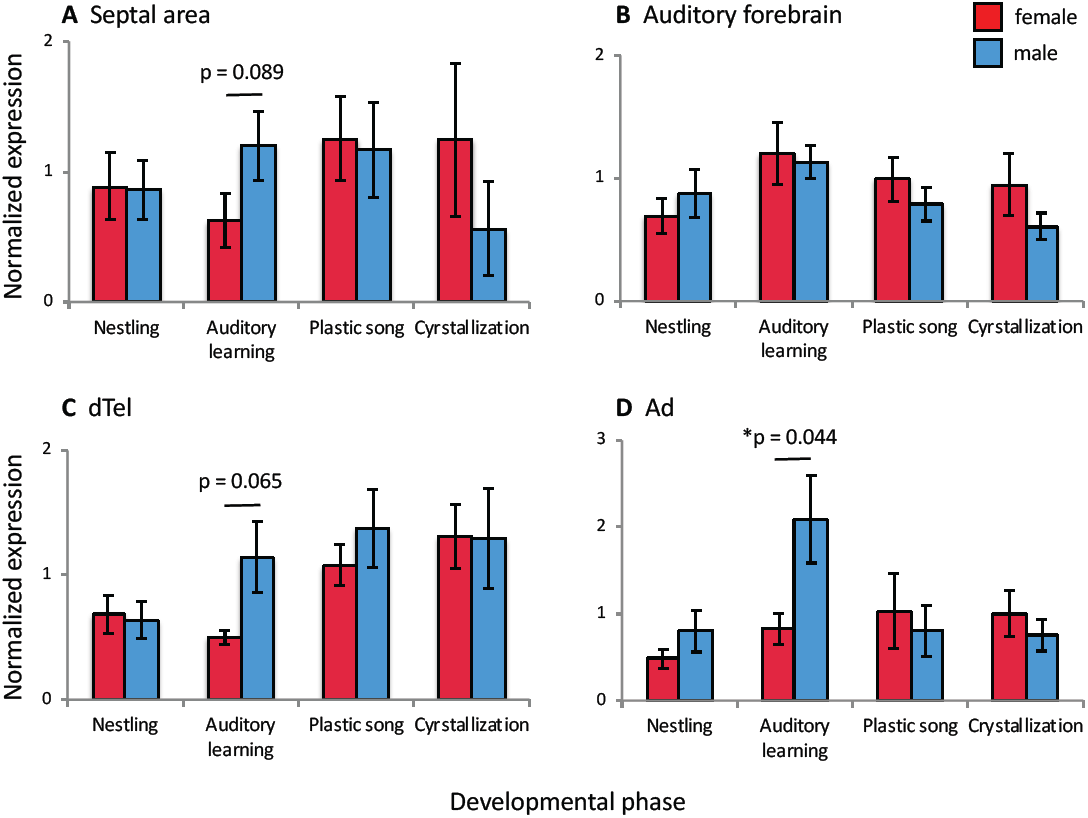
Sex differences in expression of oxytocin receptor mRNA during four phases of development. Means and SEM are shown for males (blue) and females (red) during the nestling phase (P5, P15), early auditory learning/subsong (P25, P35, P45), plastic song (P55, P65, P75) and crystallization (P85, P95). See Fig. 1 for details on how the phases were defined. Expression was normalized to two reference genes for analysis, then normalized to the series mean (within region) for graphing. Thus, 1.0 on the Y-axis corresponds to the average across both sexes and all phases for each region, and region-to-region variation is not represented here. dTel, dorsal telencephalon (includes HVC when HVC is present). Ad, dorsal arcopallium. * p < 0.05.

### Lateral septum

Some of the most intense binding of ^125^I-OVTA in adults is found in the lateral septum (Leung et al., 2009), and we found this region densely labeled in juveniles as well. We saw ^125^I-OVTA binding at high levels in the lateral septum by post-hatch day 5 (P5) (Fig. 2). The pattern of labeling was similar to what has been reported in adults (Leung et al. 2009) which was shown to overlap closely with expression of OTR mRNA (Leung et al., 2011; Fig. S1).

A univariate ANOVA showed no main effects of sex or age on OTR mRNA expression in the septal area and no interaction between sex and age (Table 1). When the age groups were organized into developmental phases (see Fig. 1), we detected a nonsignificant trend for a sex difference during auditory learning (Fig. 7A), with expression higher in males. For V1aR mRNA we detected a main effect of sex (Fig. S2A) but we found no sex difference in expression of V1aR mRNA within any of the developmental phases (Fig. S4A). For V2R mRNA there was a peak in expression in females (Fig. S3A) that appeared to drive a main effect of age and an interaction between sex and age (Table S1); however no significant sex differences were detected when birds were grouped into developmental phases (Fig. S5A).

### Auditory forebrain

In the autoradiographic films, we found dense ^125^I-OVTA labeling in both primary and secondary auditory cortex (Fig. 3; see Calabrese & Woolley, 2015 for definitions and boundaries of these regions). ^125^I-OVTA labeling was apparent in the auditory forebrain as early as P5. At all ages, this label appeared darkest in regions of Field L, primarily L1 and L3, and in the caudal nidopallium. Field L2, which receives auditory thalamic input, was unlabeled compared to surrounding regions. This pattern of dense label, surrounding but not including Field L2, was barely discernable at P5 but was obvious by P15. In adult zebra finches, this distinctive pattern of ^125^I-OVTA labeling overlaps with OTR mRNA expression more closely than it does with V1a or V2 mRNA expression (Fig. S1). The regions that are labeled respond selectively to conspecific song with the induction of immediate early genes (Mello & Clayton, 1994). The caudomedial mesopallium, which also responds selectively to song (Tomaszycki et al., 2006) and is considered part of the primary auditory cortex (Calabrese & Woolley, 2015), was not labeled by ^125^I-OVTA (Fig. 3).

In our qPCR analysis of auditory forebrain, we found a main effect of age on OTR expression that seemed to be driven by a decrease in both sexes at p55 compared with other ages (Fig. 6B). There was no main effect of sex and no interaction between sex and age (Table 1). When the age groups were organized into developmental phases (see Fig. 1), there were still no sex differences in OTR mRNA expression in the auditory forebrain at any phase (Fig. 7B). For V1aR, there was a steep decline in expression in both sexes from P5 to P15, and expression of V1aR remained low throughout the rest of the developmental period (Fig. S2B). For V2R, there were no main effects of sex or age and no interaction between them (Table S1); however when the birds were grouped into developmental phases, there was a trend for a sex difference during the plastic song phase (Fig. S5B).

### Song system

We previously observed ^125^I-OVTA binding and OTR mRNA in HVC (Leung et al., 2009; 2011; Fig. 4), which is a major song nucleus important for both learning and production of song. HVC contains neurons that are selective for tutor song in young males (Adret et al., 2012; Nick & Konishi 2005). In our previous studies in adults, labeling in HVC was highly variable; for example out of seven males, we observed OTR mRNA in HVC in only two (Leung et al., 2011). The birds in that study were adults obtained from a breeder and we did not know their ages; we had thus wondered whether the birds with clear HVC labeling might have been the youngest in that group. In the present study, we confirmed ^125^I-OVTA binding in HVC in males as young as P35 (Fig. 4). We observed signal in HVC in males, but not females, at P35 and P40. At P45 and P55 the binding in HVC was less distinct; however it became unmistakable again in males at P65 and continued to be obvious at P75, P85, and P95.

The lack of clear HVC labeling at P45 and P55 could have been related to tearing of the sections at their dorsal edge, where HVC is located. We note, however, that our qPCR data showed a reduction in OTR mRNA in males at these ages, followed by an increase by P65 (Fig. 6C). Thus it is possible that OTR expression decreases at the end of the auditory learning phase, to recover later as crystallization approaches. Although we found a main effect of sex, there was no main effect of age or interaction between sex and age (Table 1). Looking within developmental stage, we were unable to detect a significant sex difference although there was a trend during auditory learning (Fig. 7C). It is curious that, given the clear ^125^I-OVTA labeling within the boundaries of HVC in males only, we could not detect a sex difference in OTR mRNA in dTel at any age. In fact, OTR mRNA expression was remarkably similar between the sexes at P85 and P95 (Fig. 6C). This result could have been caused by generally high expression of OTR mRNA throughout the dorsal telencephalon in both sexes at the older ages. Alternatively, or in addition, our dTel punches may have sampled the periventricular region immediately overlying HVC, which expressed high levels in both sexes at all ages (see Figs. 2 and 5, and the next section of Results, below). We did not detect any effects of age or sex, or interactions, for the other two receptors in dTel (Figs S4C & S5C).

In addition to measuring OTR mRNA expression in HVC, we also looked at Ad, a region that lies immediately dorsal to a major target of HVC, the robust nucleus of the arcopallium (RA). Ad has been hypothesized to generate a comparison between memorized songs and newly generated ones, in order to influence motor output (Bottjer & Altenau, 2010). Lesions of Ad may interfere with retention of the tutor song memory over time or disrupt the ability of finches to match their own song to the songs stored in memory (Bottjer & Altenau, 2010). In adult zebra finches, Ad contains both OTR and V2 mRNA (Leung et al., 2011; Fig. S1) and concentrates ^125^I-OVTA binding (Leung et al., 2009). Here, we found clear ^125^I-OVTA labeling in Ad as seen in the autoradiographic films (Fig. 4). Although this labeling appeared to be dense in both sexes at P25, our qPCR data show a spike at that age occurring only in males (Fig. 6D). At young ages, the arcopallium labeling extended ventrally to encircle RA, in other words the RA capsular region. This labeling was seen in both sexes as early as P5 and was most pronounced around P25-35 (Fig. 4). At P55, RA capsular labeling was seen in males only, and by P75 it was weaker in both sexes while Ad remained prominently labeled.

### Periventricular hyperpallium

Perhaps the densest ^125^I-OVTA binding in the brain, at every age, was located in the periventricular region of the hyperpallium along the lateral ventricles. The distribution of this label is depicted in detail in Fig. 5, but it can be seen also in Figs. 2, 3, 4, and S1. Nearby but discontinuous labeling was also noted in an area labeled as the core of the dorsolateral corticoid area in the Zebra Finch Expression Brain Atlas (ZEBrA, zebrafinchatlas.org). Little is known about either region; both stand out as negative for expression of the immediate early gene *egr-1*, also known as *zenk*, after drug-induced induction of widespread depolarization (Mello & Clayton, 1995). The periventricular labeling includes an area known to be critical for imprinting in chicks (Aoki et al., 2015; Nakamuri et al., 2010). Early formation of social attachment to a song tutor likely involves a process similar to imprinting (Adret 1993; Baran 2017, Rodriguez-Saltos, 2017). This region may thus be important for tutor choice. Nonapeptide action in this region, and its role in social attachment, may represent an important avenue of future study.

## Discussion

In this study, we have tracked the binding of ^125^I-OVTA in both sexes of zebra finch from P5 to near-adulthood at day P95. As early as day P5, labeling was evident in the lateral septum, auditory forebrain, and song control system. These regions contained mRNA for OTR and, at lower levels, V1aR and V2R. Thus, nonapeptides have a rich set of targets on which to act throughout the period of social imprinting and vocal learning in this species, in regions that play important roles in these processes.

The caudal nidopallium for example, has received a lot of interest as a possible site of tutor song memory (Bolhuis & Moorman, 2015; Yanagihara & Yazaki-Sugiyama, 2016). In this region, sound-induced expression of *egr-1* is selective for tutor song and correlates with how well it is learned (Bolhuis et al., 2000; 2001). Song-selective *egr-1* responses develop at P35-P45 (Jin & Clayton, 1997; Stripling et al., 2001; Tomaszycki et al., 2006), which coincides with a peak in song memorization (Deshpande et al., 2014; see Fig. 1). In males reared without song tutors, conspecific song fails to elicit an *egr-1* response in the caudal nidopallium (Jin & Clayton, 1997). The region therefore appears to encode the familiarity and behavioral relevance of song (see also Maney, 2013), which may contribute to learning. Here, we showed dense ^125^I-OVTA binding in this region as well as in Field L1 and L3, which overlaps with the distribution of song-induced *egr-1* responses. Some authors have hypothesized that oxytocin may act directly within the auditory system to enhance selective responses to conspecific vocalizations (see Kanwal & Rao, 2002). Marlin et al. (2015) demonstrated in mice that oxytocin administration into the auditory cortex increased behavioral responses to pup calls, even in virgin females that would not normally respond. This result suggests that the incentive salience of pup calls was induced by changes in the cortical representation of the sound. OTR in the auditory cortex of developing zebra finches may function similarly, increasing the incentive salience of tutor song.

In all four regions in which we measured OTR mRNA, there was an interesting dip in expression in males at around P55 (Fig. 6). We observed a similarly-timed decrease in a previous study, which included juveniles at P42, P56 and P70 (Maney & Rodriguez-Saltos, 2016). At around P55, the auditory learning phase is drawing to a close (reviewed by Gobes et al., 2019) and males are entering the plastic song phase, dramatically increasing their song rate (Johnson et al., 2002). At the same time, their behavioral preferences for hearing song, as determined in a key pressing assay, shift from tutor song to other song (C. A. Rodriguez-Saltos, unpublished data). In other words, males are spending more time practicing the song they will eventually sing and less time seeking out access to the model. This is also the time at which juvenile males transition from spending time with the family unit to seeking contact with other, unrelated birds (Adkins-Regan & Leung, 2006). We hypothesize that the decrease in OTR expression at this time may mediate this shift in attention and motivation.

The capsular region surrounding song control nucleus RA was densely labeled at the younger ages (Fig. 4). Leung et al. (2009; 2011) reported that RA capsular labeling was absent in adult zebra finches; instead, the arcopallium labeling seen in adults resembled what we saw here in P75-95 juveniles. Notably, the distinctive ^125^I-OVTA labeling in capsular RA that we see here in young zebra finches was previously described in adults of several seasonally breeding species, including canaries (Voorhuis et al., 1990), house sparrows (Ondrasek et al., 2018), European starlings (Ondrasek et al., 2018), dark-eyed juncos (Wilson et al., 2016), and white-throated sparrows (Leung et al., 2009). In seasonal species, song can become highly variable, not unlike plastic song, during the non-breeding season (Smith et al., 1997); in seasonally breeding sparrows, treatment with testosterone reduced song variability in non-breeding adults (Meitzen et al., 2007) and hastened song crystallization in juveniles (Whaling et al., 1995). Treatment of male white-throated sparrows with testosterone reduced OTR mRNA in the arcopallium (Grozhik et al., 2014), suggesting that OTR in this region may be associated with song variability. Our current results are consistent with this idea in that OTR in the capsular region, as well as in the arcopallium overall, appeared to decrease in males after the auditory learning phase.

Our qPCR data showed evidence of high between-subject variation. Some of this variation, particularly for V1aR and V2R, is likely attributable to low levels of expression and high Cp values. We saw quite a bit of individual variation even in OTR mRNA expression, however. Wide variation in labeling from individual to individual is a hallmark of nonapeptide receptor expression across species (Francis et al., 2002; Leung et al., 2009; 2011; Phelps & Young, 2003). In our autoradiographic study, we had only two individuals of each sex at each age, one sectioned coronally and one sagittally. We therefore could not look for such variation in the autoradiographic films. This low sample size could explain why the effects we detected using qPCR could not always be detected on the films. For example although we detected a sex difference in OTR mRNA in Ad at around P25 (Fig. 6D), the label in the films is robust in both sexes at that age (Fig. 4). We also did not notice a decrease in signal in the auditory forebrain around P55 in the films, even though our qPCR data show this decrease both in this study (Fig. 6) and in a previous one (Maney & Rodriguez-Saltos, 2016). Given the well-documented individual variation in nonapeptide receptor expression across species, it will be necessary to conduct autoradiography on a much larger number of animals to be able to test quantitatively for changes in the signal over time.

The distribution of ^125^I-OVTA binding in our regions of interest overlaps convincingly with the distribution of OTR mRNA in adult zebra finches (Leung et al., 2011; Fig S1). This result suggests that the antagonist is labeling predominantly OTR in these regions; however, we do not know which endogenous nonapeptide is more likely to bind at these sites. Both vasotocin and mesotocin are likely to bind well to avian OTR (see Leung et al., 2011), and both nonapeptides perform oxytocin-like functions (Robinzon et al., 1994; Saito & Koike, 1992; Goodson & Bass, 2001). Our previous competitive binding studies showed that in lateral septum of adult white-throated sparrows and zebra finches, ^125^I-OVTA is displaced equally by mesotocin and vasotocin (Leung et al., 2009). In canaries, ^125^I-OVTA binding in the arcopallium is better displaced by vasotocin than by mesotocin (Voorhuis et al., 1990). Although both nonapeptides are likely being produced during the vocal learning period, studying the development of these systems in birds has been hindered by a lack of mesotocin-specific probes and antibodies that do not cross-react with vasotocin. Using a nonspecific probe to label mRNA of both nonapeptides in chick brain, Milewski et al. (1989) showed that the mRNA is detectable by embryonic day 6 and is expressed at adult-like levels in both magnocellular and parvocellular neurons by the day of hatch. Similarly, using an antibody that labels both mesotocin and vasotocin, Tennyson et al. (1986) showed that immunoreactive neurons are detectable in chick brain as early as embryonic day 6 and that the locations of mature neuronal cell groups are well-established before hatch. In canaries, vasotocin immunoreactive neurons are clearly visible in the paraventricular nucleus by P28 (Voorhuis et al., 1991). We have detected both vasotocin and mesotocin mRNA in micropunches of hypothalamus of juvenile zebra finches at P42, P56, and P70, and both mRNAs are expressed at adult-like levels at all three ages (W. M. Zinzow-Kramer, unpublished data). We are unable to detect these mRNAs in the septal area or in the auditory forebrain, which is consistent with the known distribution of nonapeptidergic cell bodies across vertebrates (Voorhuis & De Kloet, 1992; Voorhuis et al., 1991). Validation of the specificity of these probes, along with the development of specific antibodies, will be necessary to more fully understand the ontogeny of these systems in songbirds.

## Conclusions

Song learning requires attention to social stimuli. Despite the wealth of evidence that social motivation is modulated in adults of many vertebrate taxa by oxytocin, relatively little is known about how oxytocin contributes to the development of social motivation in juveniles of any species (Hammock, 2014; Miller & Caldwell, 2015). Here, we have shown that OTR mRNA is expressed in a variety of brain regions, including the auditory forebrain and song system, as early as P5 and continues to be expressed throughout song learning. If activation of these receptors facilitates attention to conspecifics, oxytocin antagonists should impair song learning. Oxytocin antagonism in the auditory cortex of mice does not block performance on a pup retrieval task if the task is already learned (Marlin et al., 2015); it should, however, block the learning of the task (Liu, 2015). Manipulating the expression or activity of nonapeptides and their receptors during song learning should prove to be a fruitful area of future research.

## Methods

### Animals

All procedures were approved by the Emory University Institutional Animal Care and Use Committee. Juvenile zebra finches were produced by a breeding colony housed in a mixed-sex aviary (12h light, 12 h dark) at Emory. All birds were free to engage in social interaction, and parents had constant access to their offspring.

We used 136 birds in total. Brains were harvested from juveniles at P5, P15, P25, P35, P45, P55, P65, P75, P85, and P95. For autoradiography, two males and two females were used for each age group; a single male at age P40 was also included. For qPCR, five males and five females were used for each age group except for P65 females (n=4) and P95 males (n=3). Birds at least 45 days old were sexed by plumage and the presence of an ovary or testis. Younger birds were sexed using PCR analysis of a small liver sample (Griffiths et al., 1998). All birds were sacrificed by isoflurane overdose and rapidly decapitated. The brains of P5 birds were left in the skull; brains of all older birds were dissected from the skulls. Brains were flash-frozen in powdered dry ice and stored at −80°C until sectioning.

### Autoradiography

For each age group we ran ^125^I-OVTA autoradiography on four brains: one of each sex in the coronal plane and one of each sex in the sagittal plane. The P40 male was sectioned on the coronal plane. Brains were sectioned on a cryostat at a thickness of 20µm and sections were thaw-mounted onto microscope slides in six parallel series. With one series of each brain, receptor autoradiography was performed using ^125^I-ornithine vasotocin (d(CH2)5[Tyr(Me)2,Thr4,Orn8,[^125^I]Tyr9-NH2]; catalog No. NEX254050UC; Perkin Elmer) as described by Leung et al. (2009). Labeled sections were exposed to Kodak BioMax maximum resolution film for seven days. The film was then developed in a Konica SRX101-A developer and scanned at 2500 dpi using a digital Epson V700 scanner. A second series of sections from each brain was Nissl-stained in order to visualize major landmarks and verify the locations of label in the autoradiographic films.

### qPCR

Brains were sectioned on a cryostat in the coronal plane. We took alternating 60µm and 300µm sections. The 60µm sections were Nissl-stained to provide a map of each 300µm section. We used the Palkovits punching technique (Palkovits & Brownstein, 1983) to extract brain tissue from the frozen 300µm sections under a dissecting microscope. All punches were taken with punch tools chilled on dry ice, and punches were immediately ejected into microtubes on dry ice. We sampled four brain regions: the septal area (targeting LS but also likely containing a portion of the medial septum), the auditory forebrain, the dorsal telencephalon (dTel) containing the HVC area, and Ad (Fig. S6). For the punches of auditory forebrain, we targeted the region that responds maximally to playback of conspecific song (see Mello & Clayton, 1994). Our sample excluded L2 but likely contained portions of L3 and the caudal nidopallium (see Vates et al., 1996). Micropunches of auditory forebrain, dTel, and Ad were taken from the same hemisphere of two adjacent sections. The hemisphere for which the borders of each region was most clearly visible in the Nissl-stained sections was selected to punch. For each tissue sample of dTel and Ad, three side-by-side 0.5mm punches were collected. A single 1mm punch was collected from the auditory forebrain. Two side-by-side 0.5mm punches were collected from the septal area in two adjacent sections, directly along the midline. Because of the small size of the P5 brains, micropunches of P5 birds were obtained using slightly different methods, as follows: for the septal area, a single 0.5mm punch was obtained directly on the midline in two adjacent sections. For the auditory forebrain, a single 1mm punch was obtained from each hemisphere. dTel, Ad, and the auditory forebrain were sampled bilaterally in one section only. For dTel and Ad in P5 birds, two side-by-side 0.5mm punches were obtained from each hemisphere. All punches were stored at −80°C for a maximum of 2 days before RNA isolation.

We extracted RNA from each punch using the Qiagen Allprep RNA/DNA micro kit with modifications (Zinzow-Kramer et al., 2015). To produce cDNA, we performed reverse transcription using the Transcriptor First Strand cDNA synthesis kit with random hexamer primers, then diluted the reaction to 20 ng/µl for qPCR. We designed exon-spanning primers (Table S2) for use with probes from the Roche Universal Probe Library for OTR, V1aR, and V2R, then verified the amplified sequences via cloning and sequencing. qPCR was performed using a Roche LightCycler 480 Real-Time PCR System, in triplicate for each sample on a 384-well plate as previously described (Zinzow-Kramer et al., 2014). The target genes (OTR, V1aR, and V2R) and reference genes were always run on the same plate for each sample, and sex and age were balanced across plates. Using the LightCycler 480 Software Version 1.5.0, we calculated crossing point values using the Abs Quant/2nd Derivative Max method. We normalized the expression of each gene of interest to the geometric mean of two reference genes (Pfaffl, 2001; Vandesompele et al., 2002). The reference genes were glyceraldehyde 3-phosphate dehydrogenase (GADPH) and peptidylprolyl isomerase A (PPIA), which have been previously validated for use in zebra finch brain tissue and are stably expressed across sex (Zinzow-Kramer et al., 2014). For samples with a coefficient of variance above 21.8% within triplicates, indicating technical error, we identified and removed one outlying point within the triplicate (Bookout et al., 2006) using a modified Dixon’s q test (Dean & Dixon, 1951).

### Statistical analyses of gene expression

For the data across all animals within each gene and region, we removed outliers using SPSS v26, which uses the interquartile range with 3.0 as a multiplier. We then analyzed the entire dataset in an omnibus repeated-measures MANOVA with gene and region as repeated measures and sex, age, and qPCR plate as between-subjects factors. For the MANOVA, the missing values (removed outliers) were replaced with the series mean. Because we found a significant gene x region x sex x age interaction in this MANOVA (see Results), we then performed univariate ANOVAs for each gene within each region, with sex, age, and qPCR plate as between-subjects factors. Missing values were not replaced for these univariate tests.

Due to limited power, we did not test for sex differences within each of the ten age groups. To increase our power to detect sex differences, we lumped age groups together into developmental phases. Our groupings (see Fig. 1 for our rationale) were “nestling” (P5, P15), “auditory learning” (P25, P35, P45), “plastic song” (P55, pP65, P75) and “crystallization” (P85, P95). We then ran an omnibus repeated-measures MANOVA as before, with gene and region as the repeated measures and sex, phase, and qPCR plate as between-subjects factors. Upon finding a gene x region x sex x phase interaction (see Results) we then ran univariate ANOVAs within gene and region. These ANOVAs were performed within each phase, testing for a main effect of sex while controlling for qPCR plate.

## Author Contributions and Notes

MTD, KEG, and DLM designed the studies, MTD and KEG performed the procedures, MTD and DLM analyzed the data, and MTD and DLM wrote the paper.

The authors declare no conflict of interest.

This article contains supporting information online.

## Acknowledgments

We are grateful to T. J. Libecap, Kiyoshi Inoue, Cary Leung, Jennifer Merritt, and Wendy Zinzow-Kramer for technical assistance, and to Sarah London for help with nestling neuroanatomy. We also thank Erich Jarvis and Laura Carruth for providing founders for our zebra finch colony. This work was supported by NIH 1R21MH105811-01A1.

