## Supplemental Tables and Figures for "Expression of oxytocin receptors in the zebra finch brain during vocal development"

**Table S1.** Results of univariate F-tests within gene and region, showing main effects of age (P5 – P95) and sex on expression of the V1a and V2 receptors, and interactions between age and sex, in the four regions of interest. Significant effects shown in bold and marked by asterisks.

| | | **Age** | | **Sex** | | **Age x Sex** | |
|---|---|---|---|---|---|---|---|
| **Gene** | **Region** | **F** | ***p*** | **F** | ***p*** | **F** | ***p*** |
| **V1aR** | Septum | 1.782 | 0.150 | **9.575** | **0.007*** | 1.904 | 0.125 |
| | NCM | **3.304** | **0.015*** | 0.180 | 0.677 | 0.752 | 0.660 |
| | Dorsal telencephalon | 1.367 | 0.277 | 0.784 | 0.388 | 0.185 | 0.993 |
| | Ad | 0.476 | 0.874 | 1.657 | 0.213 | 0.614 | 0.771 |
| **V2R** | Septum | **13.792** | **<0.001*** | 1.089 | 0.321 | **9.731** | **0.001*** |
| | NCM | 1.042 | 0.451 | 0.772 | 0.392 | 1.016 | 0.467 |
| | Dorsal telencephalon | 2.004 | 0.108 | 0.123 | 0.731 | 2.415 | 0.063 |
| | Ad | 1.468 | 0.246 | 0.076 | 0.787 | 1.852 | 0.140 |

**Table S2**. Reference and target genes quantified with qPCR.

| **Gene** | **Forward primer** | **Reverse primer** | **Roche probe no.** | **Gene accession no.** |
|---|---|---|---|---|
| GAPDH | CAACTTCGGCATTGTGGAG | GGCCCATCCACTGTCTTCT | 76 | NM_001198610.1 |
| PPIA | GAAGGGCTTCGGCTACAAG | CCATTGTGGCGTGTGAAGT | 38 | NM_001245462.1 |
| OTR | GATGTGGTCTGTGTGGGACA | TTGCAGCAGCTGTTGAGG | 4 | JN594029.1 |
| V1aR | TCATCGTCTCGGTGTACGTC | AGTTGCAGTGTTTTCAGAATCG | 107 | JN594032.1 |
| V2R | CACCTTTCTTCATTGCACAGC | TAATGGTGAATGCCGAACCT | 89 | JN594025.1 |

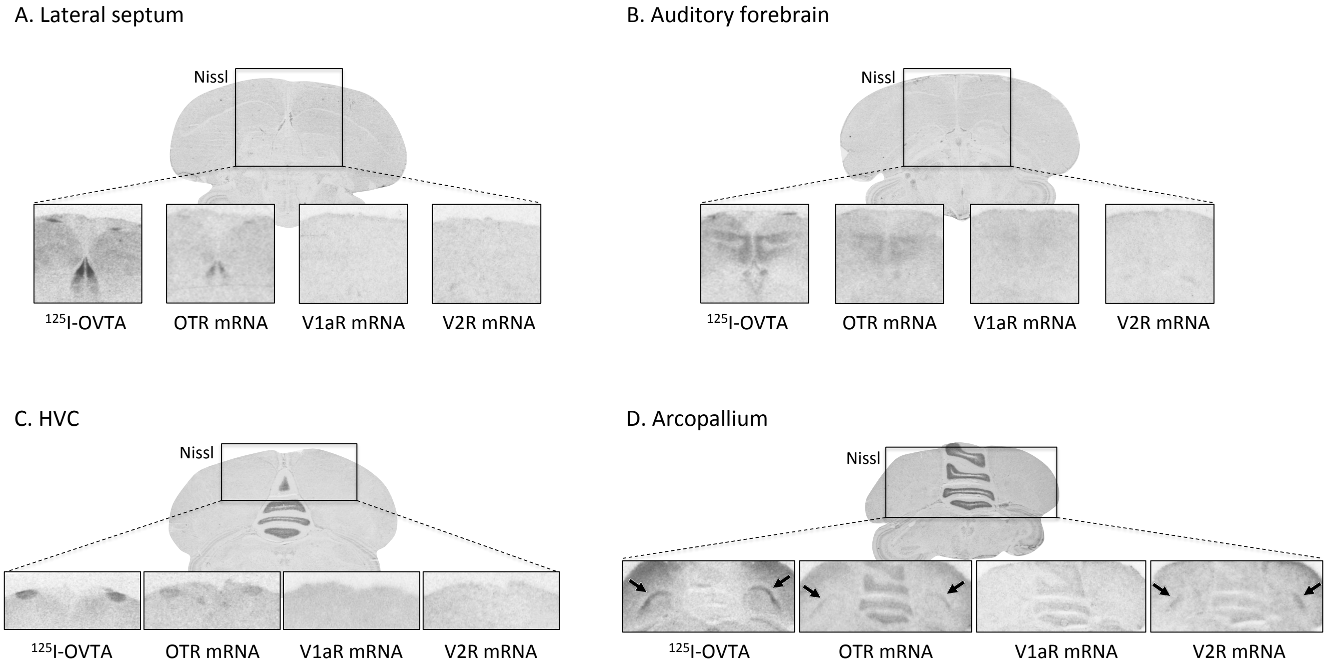


**Fig. S1. The distribution of $^{125}$I-OVTA binding closely matches the distribution of oxytocin receptor (OTR) mRNA in four regions of the zebra finch brain.** In A-D, $^{125}$I-OVTA labeling (autoradiography) is compared with expression of OTR, V1aR, and V2 mRNAs (*in situ* hybridization) in adjacent coronal sections from the same animal. The Nissl-stained section above each set of photos is the same section in which $^{125}$I-OVTA was labeled. (A) the distribution of $^{125}$I-OVTA labeling in lateral septum resembles OTR mRNA expression more closely than it resembles V1aR mRNA or V2 mRNA. Some signal is also present for V1aR, but it is much weaker. Note labeling for $^{125}$I-OVTA and OTR mRNA not only in the lateral septum but also in the periventricular area. See also Fig. 2 in main text. (B) $^{125}$I-OVTA binds to receptors in the auditory forebrain in a pattern similar to the distribution of OTR mRNA. Field L2 is relatively unlabeled compared with the surrounding caudomedial nidopallium and other Field L subregions. See also Fig. 3 in main text. (C) $^{125}$I-OVTA distinctly labels song control nucleus HVC, in a pattern similar to that of OTR mRNA. See also Fig. 4 in the main text. (D) $^{125}$I-OVTA binding is seen in the dorsal arcopallium (arrows), which matches the pattern of OTR mRNA. See also Fig. 4 in the main text. Note, however, that V2R mRNA can also be detected in this region (arrows in rightmost panel). All material is from Leung et al. (2009; 2011). A-C are from the same adult male. D is from an adult female.

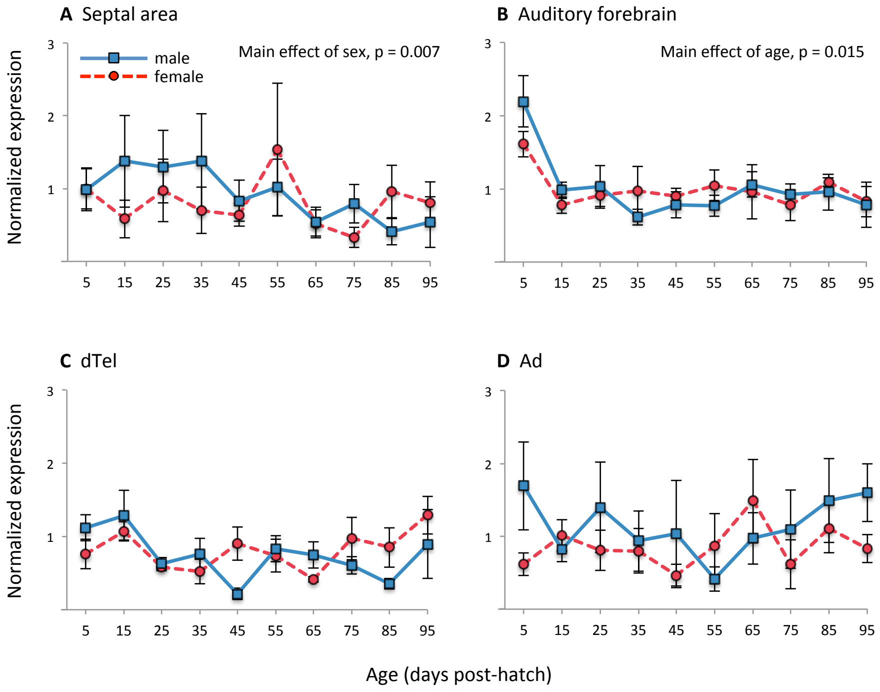


**Fig. S2. Developmental trajectories of V1aR expression in four brain regions of interest.** Means and SEM are shown for males (blue) and females (red) at ten ages post-hatch. Expression was normalized to two reference genes for analysis, then normalized to the series mean (within region) for the purposes of graphing. Thus, 1.0 on the Y-axis corresponds to the average across both sexes and all ages for each region, and region-to-region variation is not represented here. dTel, dorsal telencephalon (includes HVC when HVC is present). Ad, dorsal arcopallium.

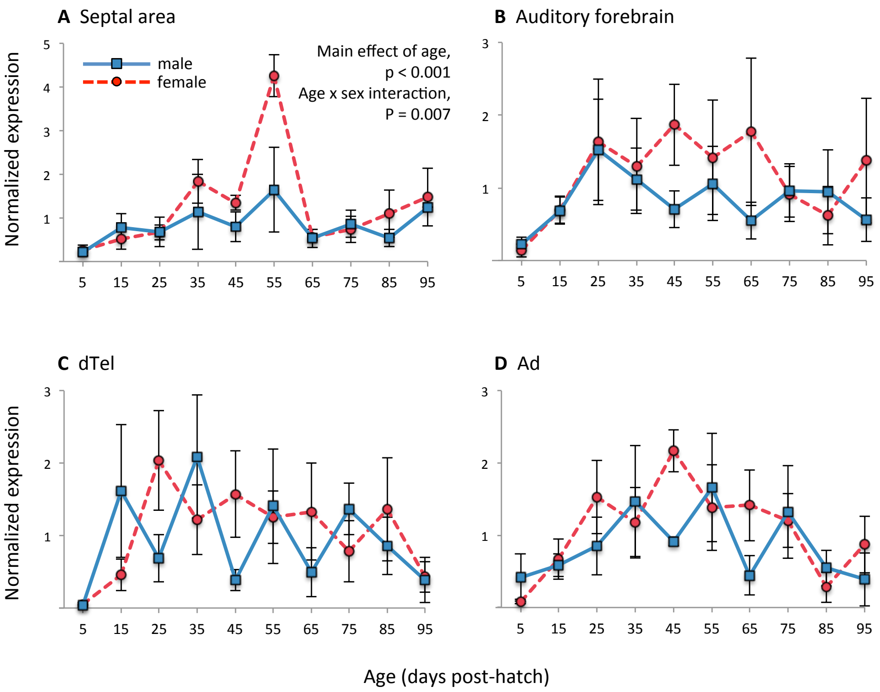


**Fig. S3. Developmental trajectories of V2R expression in four brain regions of interest.** Means and SEM are shown for males (blue) and females (red) at ten ages post-hatch. Expression was normalized to two reference genes for analysis, then normalized to the series mean (within region) for the purposes of graphing. Thus, 1.0 on the Y-axis corresponds to the average across both sexes and all ages for each region, and region-to-region variation is not represented here. dTel, dorsal telencephalon (includes HVC when HVC is present). Ad, dorsal arcopallium.

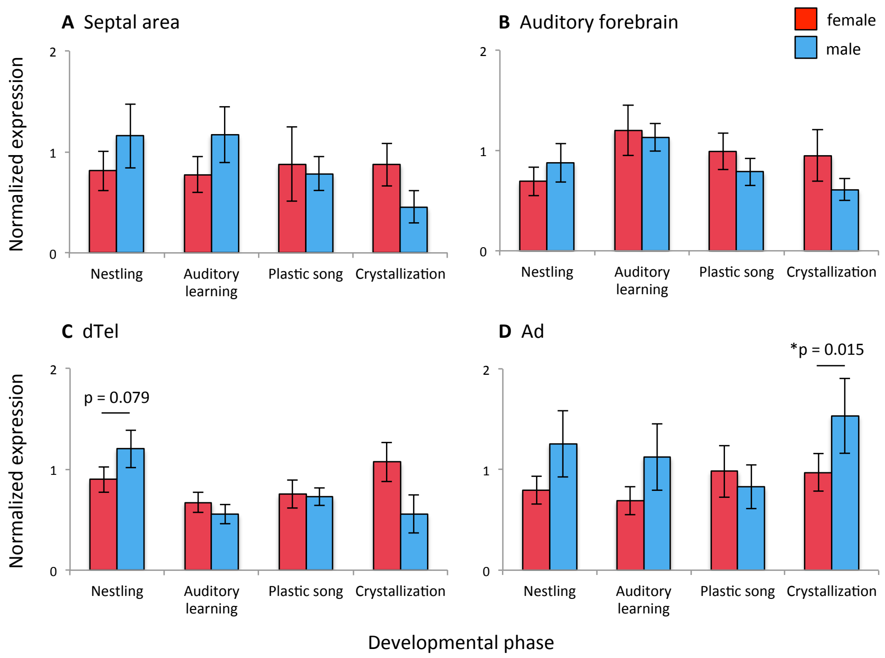


**Fig. S4. Sex differences in V1aR expression during four phases of development.** Means and SEM are shown for males (blue) and females (red) during the nestling phase (P5, P15), early auditory learning/subsong (P25, P35, P45), plastic song (P55, P65, P75) and crystallization (P85, P95). See Fig. 1 for details on how the stages were defined. Expression was normalized to two reference genes for analysis, then normalized to the series mean (within region) for the purposes of graphing. Thus, 1.0 on the Y-axis corresponds to the average across both sexes and all phases for each region, and region-to-region variation is not represented here. dTel, dorsal telencephalon (includes HVC when present). Ad, dorsal arcopallium. *$p < 0.05$.

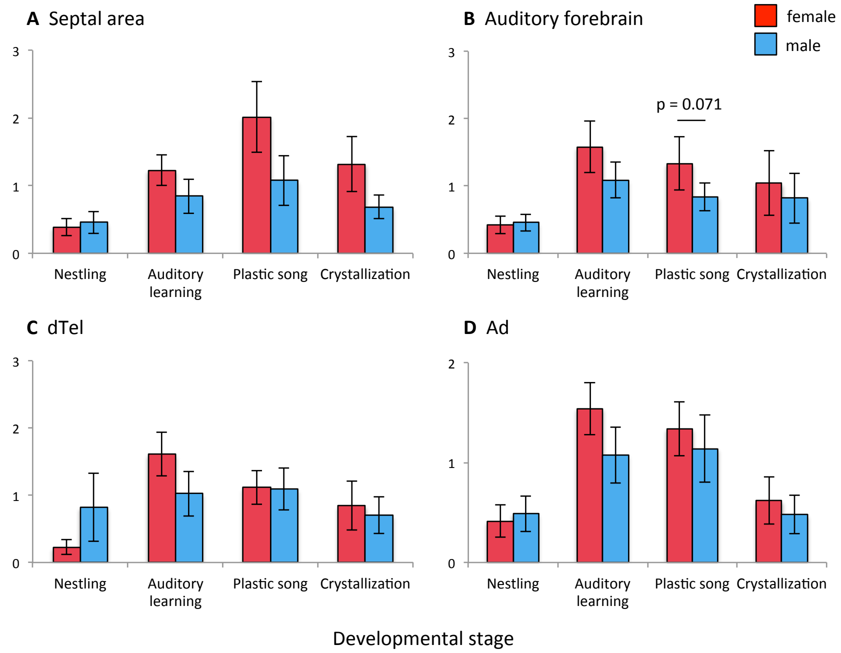


**Fig. S5. Sex differences in V2R expression during four phases of development.** Means and SEM are shown for males (blue) and females (red) during the nestling phase (P5, P15), early auditory learning/subsong (P25, P35, P45), plastic song (P55, P65, P75) and crystallization (P85, P95). See Fig. 1 for details on how the stages were defined. Expression was normalized to two reference genes for analysis, then normalized to the series mean (within region) for the purposes of graphing. Thus, 1.0 on the Y-axis corresponds to the average across both sexes and all phases for each region, and region-to-region variation is not represented here. dTel, dorsal telencephalon (includes HVC when present). Ad, dorsal arcopallium. *$p < 0.05$.

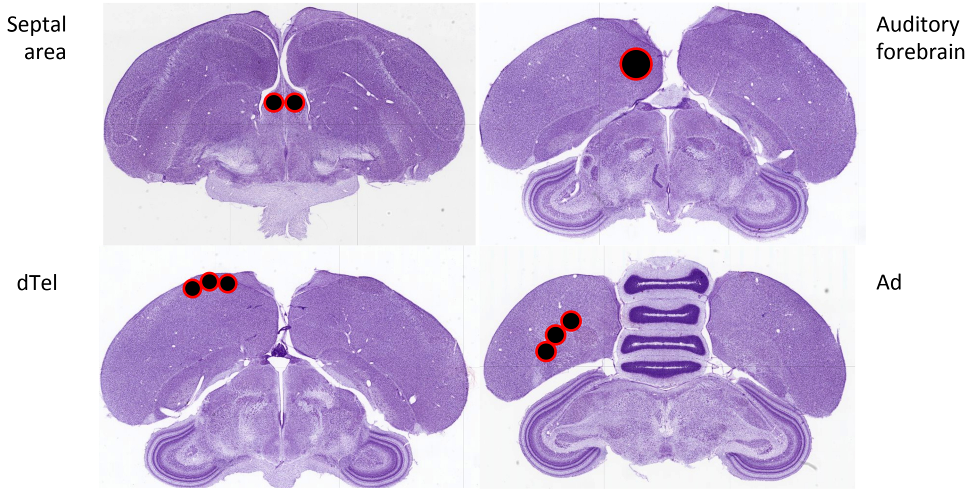


**Fig. S6. Locations of micropunches for quantitative PCR.** Above, Nissl-stained sections are shown to indicate the locations of our punches of fresh frozen 300μm sections. The punch of auditory forebrain was 1mm; other punches were 0.5mm. Ad, dorsal arcopallium. dTel, dorsal telencephalon.
